# Mapping Isoform Abundance and Interactome of the Endogenous TMPRSS2-ERG Fusion Protein with Orthogonal Immunoprecipitation-Mass Spectrometry Assays

**DOI:** 10.1101/2020.09.23.309088

**Authors:** Zhiqiang Fu, Yasmine Rais, X. Chris Le, Andrei P. Drabovich

**Author notes:** Correspondence to: A. P. Drabovich, Ph.D., Department of Laboratory Medicine and Pathology, University of Alberta, 10-102 Clinical Sciences Building, Edmonton, Alberta, Canada T6G 2G3.

## Abstract

TMPRSS2-ERG gene fusion, a molecular alteration driving nearly a half of prostate cancer cases, has been intensively characterized at the transcript level, while limited studies explored the molecular identity and function of the endogenous fusion at the protein level. Here, we developed and applied immunoprecipitation-mass spectrometry (IP-MS) assays for the measurement of a low-abundance T1E4 TMPRSS2-ERG fusion protein, its isoforms and its interactome in VCaP prostate cancer cells. IP-MS assays quantified total ERG (∼27,000 copies/cell) and its four unique isoforms, and revealed that the T1E4-ERG isoform accounts for 71% of the total ERG protein in VCaP cells. For the first time, the N-terminal peptide (methionine-truncated and N-acetylated TASSSSDYGQTSK) unique for the T1/E4 fusion was identified and quantified. IP-MS with the C-terminal antibodies identified 29 proteins in the ERG interactome, including SWI/SNF chromatin remodeling complex subunits and numerous transcriptional co-regulators. Our data also suggested that TMPRSS2-ERG protein-protein interactions were exerted through at least two different regions. Knowledge on the distinct TMPRSS2-ERG protein isoforms and interactomes may facilitate development of more accurate diagnostics and targeted therapeutics of prostate cancer.

## Introduction

Prostate cancer is the most frequently diagnosed neoplasm and the third leading cause of cancer mortality in men. Introduction of prostate specific antigen (PSA) testing revolutionized the practice of urologic oncology,^1^ facilitated earlier detection of localized tumors, and resulted in the active surveillance as a treatment option for many patients with the low-grade prostate cancer. PSA test, however, is prone to over-diagnosis and fails to differentiate between indolent and aggressive cancers.^2,3^ The race for prognostic biomarkers continues, with numerous genomic, transcriptomic, proteomic and metabolomic markers being recently discovered and validated.^4,5^ The most promising biomarkers are also explored for the molecular mechanisms of their differential expression or regulation.^6-8^

Recent genomic studies on the primary prostate adenocarcinoma revealed major subtypes defined by the gene fusions of E26 transformation-specific (ETS) transcription factors, and mutations in SPOP, FOXA1, and IDH1 genes.^9,10^ The most common genomic subtype of primary prostate cancer was represented by the fusion of the androgen-responsive gene TMPRSS2 with the transcription factor ERG (∼50% of all cases).^11^ While TMPRSS2-ERG rearrangement is heterogeneous, the fusion of TMPRSS2 exon 1 with the ERG exon 4 (T1/E4) occurs in ∼80% of all TMPRSS2-ERG cases.^12^ Functionally, TMPRSS2-ERG fusion results in the androgen-dependent over-expression of the N-terminally truncated ERG protein and its isoforms, which initiate and drive oncogenic transformation of the prostate epithelial cells (**Figure 1A**).^13^

**Figure 1.**
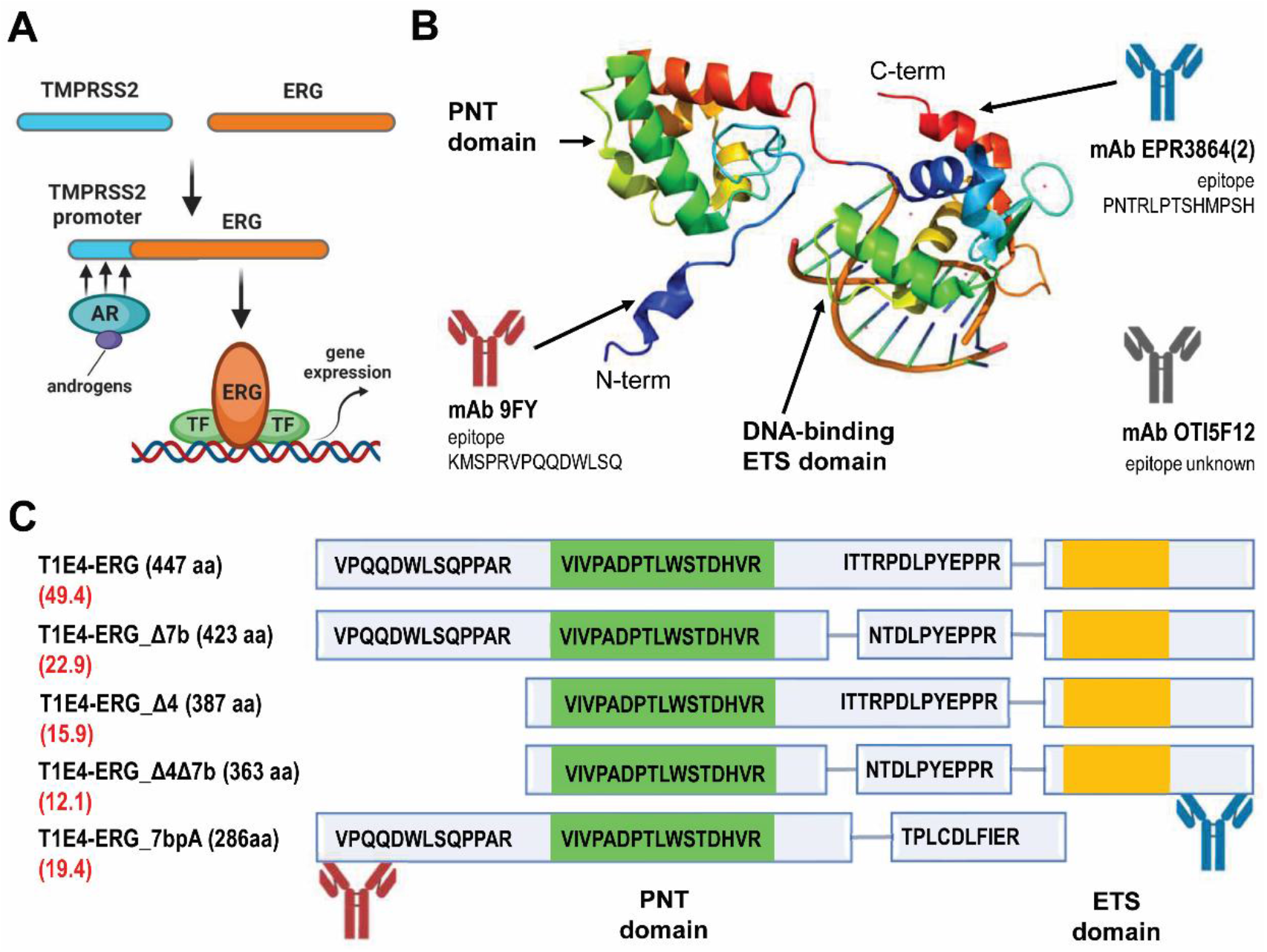
Experimental setup. (A) Androgen receptor (AR)-mediated over-expression of the TMPRSS2-ERG fusion promotes, in coordination with other transcription factors (TF), oncogenic transformation of prostate epithelial cells. (B) Schematic illustration of ERG protein structure from an arbitrary combination of the N-term (PDB ID: 1SXE) and C-term PDB structures (PDB ID: 4IRI), and corresponding positions of mAb epitopes. 5F12 mAb epitope is not known, but is likely an internal epitope; (C) Location of unique and shared peptides within different ERG isoforms (transcripts per million abundance of each mRNA isoform is shown in red; CCLE data); the PNT (Pointed) and DNA-binding ETS domains are highlighted in green and yellow, respectively.

Numerous studies evaluated TMPRSS2-ERG fusion mRNA as a prognostic biomarker, but paradoxically revealed positive, negative, or no association with the PCa clinical significance, progression or aggressiveness.^14, 15^ At the protein level, quantification of ERG has been suggested as a surrogate marker for the TMPRSS2-ERG fusion. Measurement of ERG protein expression by immunohistochemistry (IHC) in the prostate needle biopsies^16,17^ was proposed to diagnose limited adenocarcinoma, resolve atypical glands suspicious for adenocarcinoma, detect pre-cancerous lesions, or select patients for targeted therapeutic interventions.^18^ Antibody-based IHC analysis, however, could not provide any details on the heterogeneity of ERG protein isoforms or post-translational modifications.

ERG gene has twelve predicted protein-coding mRNA isoforms, of which 6 isoforms were experimentally detected by RNA sequencing in VCaP cells (**Supplemental Table S1**). While the expression of the wild-type ERG protein in VCaP cells is negligible, T1/E4 fusion results in over-expression of the N-term truncated isoforms which retain the function the full ERG. Earlier studies suggested that some ERG isoforms could have distinct molecular functions.^19, 20^ For instance, high levels of a presumably protein-coding mRNA isoform-8 were frequently detected in VCaP cells and patient tissues.^12^ Due to the missing DNA-binding ETS domain, isoform-8 protein was suggested to interfere with the transcriptional activation mediated by the full-length ERG isoforms. Recombinant isoform-8 was used as a proxy to investigate its molecular function,^20^ but the expression of the endogenous isoform-8 protein has never been demonstrated in VCaP cells or prostate tissues. Apart from ERG isoforms, there is still limited experimental data available for the post-translational modifications,^21^ protein domains, and interactomes^22-24^ of the endogenous ERG protein.

Quantitative proteomic by mass spectrometry^25-28^ is a promising tool to generate novel knowledge on TMPRSS2-ERG heterogeneity at the protein level. He et al. pioneered measurements of ERG protein by targeted mass spectrometry, utilizing two-dimensional liquid chromatography separations and selected reaction monitoring (SRM) assays.^29, 30^ Unique peptides of the total ERG protein were quantified in VCaP cell lysate, achieving limits of detection of 1.8 fg/cell or ∼3,000 cells spiked into urine.

In this study, we aimed at developing immunoprecipitation-shotgun mass spectrometry (IP-MS) and immunoprecipitation-selected reaction monitoring (IP-SRM) assays for the identification and quantification of the endogenous low-abundance TMPRSS2-ERG fusion protein and its isoforms. As a model cell line, we selected VCaP prostate cancer epithelial cells which harbored T1/E4 TMPRSS2-ERG gene fusion, over-expressed T1/E4 fusion mRNA, and expressed detectable levels of the endogenous ERG protein. We hypothesized that our sensitive IP-MS and IP-SRM assays, as well as orthogonal assays designed with the N-term and C-term monoclonal antibodies, will provide novel knowledge on the abundance of the endogenous ERG isoforms, their post-translational modifications, and ERG protein-protein interactions.

## Results

### Selection of ERG protein isoforms

In order to quantify the expression and relative abundance of different ERG isoforms, tryptic peptides for the total ERG, peptides shared by some isoforms, or peptides unique for specific isoforms were selected. RNA sequencing data available at the Cancer Cell Line Encyclopedia (CCLE) and DeepMap database (v2; https://depmap.org^31^) revealed that only 6 protein-coding mRNA isoforms were expressed in VCaP cells (**Table S1**). Our rationale for consideration of endogenous ERG isoforms was based on the following facts: (i) VCaP cells have four copies of chromosome 21,^32^ of which only two harbor T1/E4 TMPRSS2-ERG fusion and express TMPRSS2-ERG mRNA and protein; (ii) wildtype chromosomes 21 do not express any wild-type ERG (**Table S4, Figure S2**), similar to the CCLE prostate cancer cells without this fusion; (iii) quantification of mRNA isoform expression by CCLE, based on the RNA-Seq by Expectation Maximization (RSEM) algorithm, does not consider the lack of the initial three exons of ERG in VCaP cells. As a result, the assignment of isoform abundances may not be correct for some pairs of isoforms (for example, isoforms 3 and 5). In addition, mRNA T1/E4 isoforms 3 and 5, as well as isoforms 2 and B5MDW0, are identical at the protein level (**Table 1**); (iv) theoretical protein-coding mRNA isoforms 7, A0A088AWP2, A8MX39, A8MZ24 have unique internal exons, so the lack of their expression in VCaP cells based on RSEM data is apparent. As a result, we considered for further investigation five protein isoforms (**Table 1**).

**Table 1.**
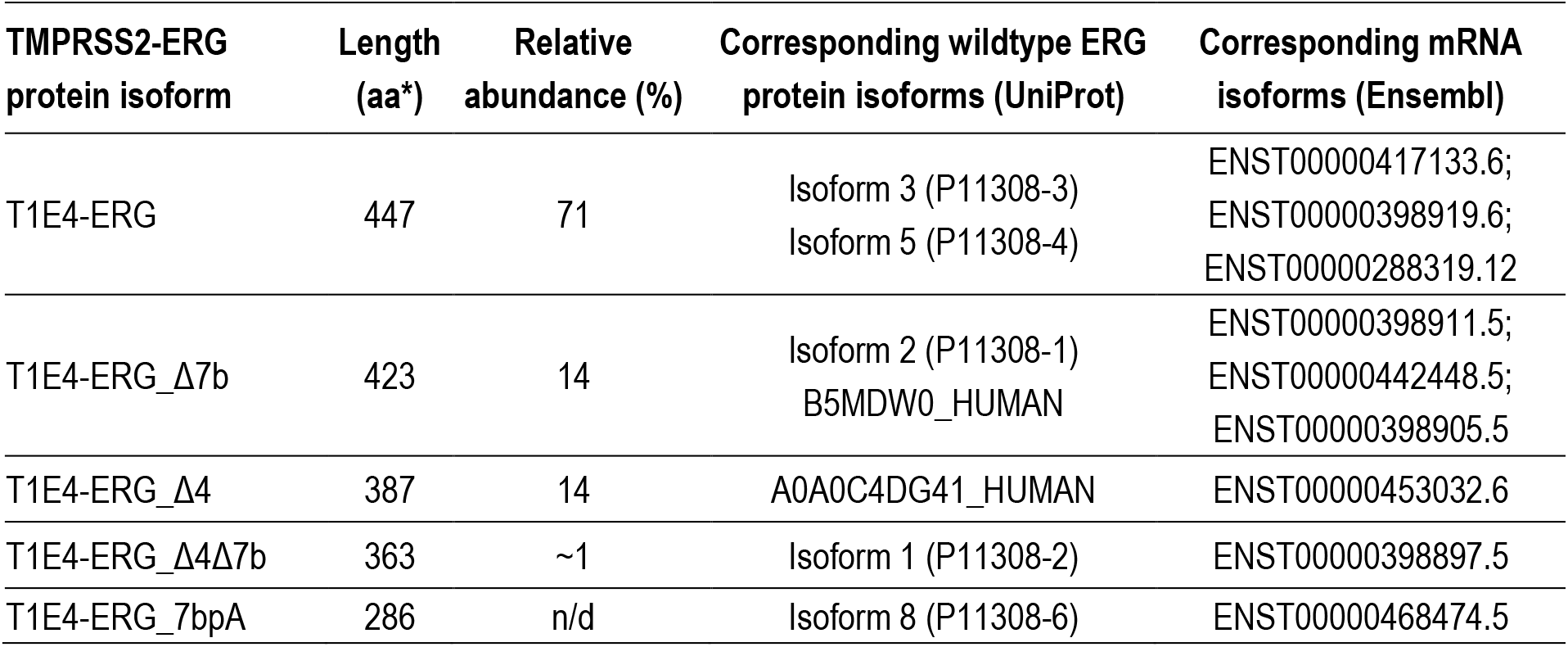
TMPRSS2-ERG protein isoforms identified and quantified in VCaP cells. Protein isoform naming is according to Zammarchi et al.^58^ T1E4, fusion of TMPRSS2 exon 1 with ERG exon 4; Δ4 and Δ7b, isoforms arising due to the missing exons 4 or 7b; 7bpA, an isoform arising due to the alternative polyadenylation site; n/d – not detected. *Protein length includes the N-terminal methionine.

| TMPRSS2-ERG protein isoform | Length (aa*) | Relative abundance (%) | Corresponding wildtype ERG protein isoforms (UniProt) | Corresponding mRNA isoforms (Ensembl) |
| --- | --- | --- | --- | --- |
| T1E4-ERG | 447 | 71 | Isoform 3 (P11308-3)<br>Isoform 5 (P11308-4) | ENST00000417133.6;<br>ENST00000398919.6;<br>ENST00000288319.12 |
| T1E4-ERG_Δ7b | 423 | 14 | Isoform 2 (P11308-1)<br>B5MDW0_HUMAN | ENST00000398911.5;<br>ENST00000442448.5;<br>ENST00000398905.5 |
| T1E4-ERG_Δ4 | 387 | 14 | A0A0C4DG41_HUMAN | ENST00000453032.6 |
| T1E4-ERG_Δ4Δ7b | 363 | ~1 | Isoform 1 (P11308-2) | ENST00000398897.5 |
| T1E4-ERG_7bpA | 286 | n/d | Isoform 8 (P11308-6) | ENST00000468474.5 |

Among those five protein isoforms, only T1E4-ERG_7bpA had unique peptides (**Figure 1C**). The remaining four isoforms could be distinguished using combinations of “razor” tryptic peptides, such as VPQQDWLSQPPAR, NTDLPYEPPR, and ITTRPDLPYEPPR (**Figure 1C**). Since isoforms T1E4-ERG_Δ4Δ7b and T1E4-ERG_Δ4 are lacking the N-term exon 4 with an epitope for the 9FY monoclonal antibody (KMSPRVPQQDWLSQ), immunoprecipitation with either the N-term 9FY antibody or C-term EPR3864(2) antibody (epitope PNTRLPTSHMPSH) followed by quantification of unique, razor and total ERG peptides facilitated differential quantification of the relative abundances of these distinct ERG isoforms.

### Development of orthogonal IP-SRM assays

Heavy isotope-labeled peptides were monitored with the multiplexed SRM assay. Based on the SRM peak area, the dominant charge states and collision energies for each peptide were determined, and poorly performing peptides were removed. Transitions scan times were optimized in the matrix of VCaP cell lysates, and peptides were scheduled within 2-min acquisition windows (**Table S5)**. For correct identification of endogenous peptides, superposition of peaks of the heavy standards and light peptides, identical peak shapes, and the same order of peak heights for heavy and light transitions were ensured. The abundance of light endogenous peptides was determined by the peak area ratio of light peptide to a heavy peptide standard spiked into each sample. The peak area was calculated as the integrated area of all transitions for each peptide.

### Quantification of TMPRSS2-ERG protein isoforms by orthogonal IP-SRM assays

ERG protein was immunoprecipitated from VCaP cell lysate with three different anti-ERG monoclonal antibodies coated on a 96-well plate. Following trypsin digestion and proteomic sample preparation, ERG protein was quantified by SRM using trypsin-cleavable SpikeTides_TQL internal standard peptides (**Table S6**), to enable “absolute” quantification (**Figure 2A**). **Figure 2B** shows the extracted-ion chromatograms of transitions monitored for unique ERG peptides detected in VCaP lysates. The amounts of ERG peptides detected in VCaP lysates by three different antibodies are listed in **Table 2**, with detailed peak areas listed in **Table S7**. The recovery of ERG after IP was estimated at ∼90% (as compared to the total ERG amount measured in VCaP digest without IP). IP-SRM assays revealed excellent reproducibility, with coefficient of variation (CV) below 10% across all samples (**Table 2**). Limit of detection of IP-SRM, as estimated with serial dilutions of VCaP lysate, was 93.6 amol/µg total protein or ∼10,000 cells **(Figure S3)**. Three antibodies displayed similar efficiency of capturing total ERG protein (69-84%), as revealed by the enrichment versus isotype controls (**Figure 2C and Table S8**). Based on the cell count, the amount of total ERG was estimated as 2.2 fg per VCaP cell (∼27,000 copies/cell), which was in good agreement with the previously reported levels (1.8 fg per cell).^29^

**Table 2.** Quantification of TMPRSS2-ERG isoform razor peptides by IP-SRM in VCaP cells. The amounts and coefficients of variation (CV) were calculated based on analytical triplicates, with detailed data presented in **Table S7**.

| Shared and unique peptides | ERG isoforms | N-term mAb;<br>fmols (CV%) | C-term mAb;<br>fmols (CV%) | 5F12 mAb;<br>fmols (CV%) |
| --- | --- | --- | --- | --- |
| VIVPADPTLWSTDHVR | Total ERG (all 5 isoforms) | 0.015 (8.4) | 0.013 (2.2) | 0.012 (0.3) |
| VPQQDWLSQPPAR | T1E4-ERG; T1E4-ERG_Δ7b;<br>T1E4-ERG_7bpA | 0.019 (6.2) | 0.011 (6.0) | 0.009 (1.7) |
| ITTRPDLPYEPPR | T1E4-ERG; T1E4-ERG_Δ4 | 0.004 (6.4) | 0.004 (9.1) | 0.003 (5.0) |
| NTDLPYEPPR | T1E4-ERG_Δ7b;<br>T1E4-ERG_Δ4Δ7b | 0.001 (7.0) | 0.001 (7.0) | 0.001 (2.1) |
| TPLCDLFIER | T1E4-ERG_7bpA | 0 | 0 | 0 |

**Figure 2.**
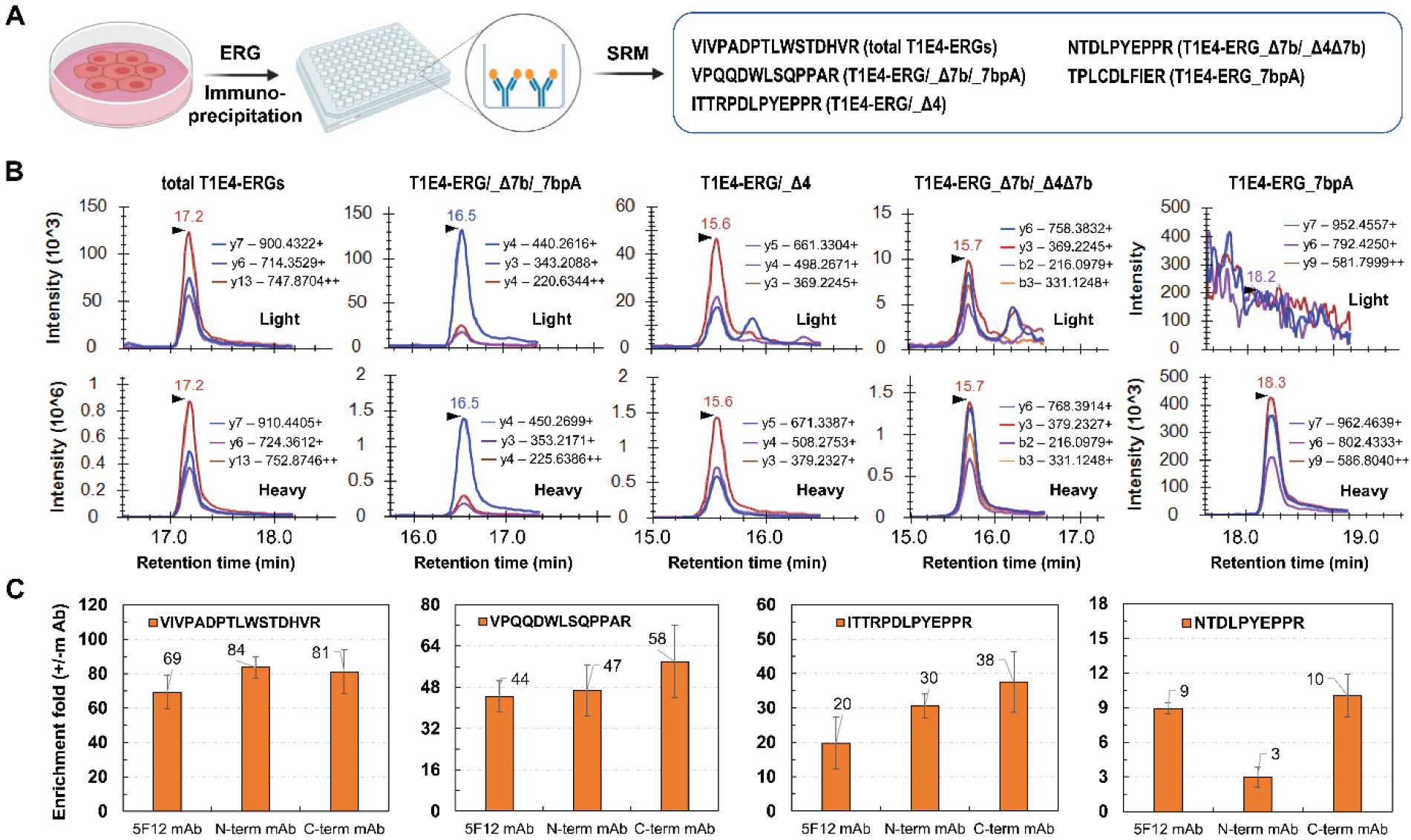
IP-SRM measurement of shared and unique ERG isoform peptides in VCaP cells. (A) Schematic workflow of the IP-SRM experiments and sequences of targeted peptides; (B) SRM measurement of endogenous ERG peptides; (C) Enrichment fold changes ((+)mAb/(-)mAb peak area ratios) revealed that three mAbs effectively captured the total T1E4-ERGs from the VCaP lysate, while the N-term mAb failed to capture the T1E4-ERG_Δ4 and T1E4-ERG _ Δ4Δ7b isoforms lacking the N-term epitope.

None of the unique peptides of T1E4-ERG_7bpA protein isoform (mRNA isoform-8) were detected (**Figure S4**), while detection of mutually exclusive peptides ITTRPDLPYEPPR and NTDLPYEPPR indicated expression of at least two distinct ERG isoforms (**Figure 2B)**. Based on our data (**Tables 1** and **2)**, the relative abundance of four ERG isoforms was estimated (see details in the **Supplemental text**) as 71% (T1E4-ERG), 14% (T1E4-ERG_Δ7b), 14% (T1E4-ERG_Δ4), and 1% (T1E4-ERG_Δ4Δ7b). To the best of our knowledge, the relative abundance of endogenous ERG isoforms, as well as the lack of expression of the shortest isoform T1E4-ERG_7bpA lacking the ETS domain (mRNA isoform-8) was quantified for the first time.

### Identification of the N-terminal peptide of T1/E4 TMPRSS2-ERG protein

Tryptic digestion of the two most abundant protein isoforms (T1E4-ERG and T1E4-ERG_Δ7b; 85% of the total ERG in VCaP cells) should generate a unique N-terminal peptide MTASSSSDYGQTSK, while the corresponding wild-type ERG isoforms would generate a longer tryptic peptide TEMTASSSSDYGQTSK. However, previous studies,^29, 30^ as well as our earlier attempts, could not detect the endogenous MTASSSSDYGQTSK peptide.

We thus assumed that the N-terminus of ERG could be further modified through acetylation, methionine cleavage, and threonine or serine phosphorylation. Our motivation to reveal the N-terminal identity of ERG stemmed not only from the challenge of detecting a unique fusion peptide, but also from the fact the knowledge on the N-term structure of ERG protein could be useful to elucidate the increased stability of TMPRSS2-ERG fusion protein,^33^ and facilitate development of the targeted therapeutic strategies for inhibition or degradation^34^ of oncogenic, but not the wild-type ERG forms.

To reveal the N-terminal identity of the T1/E4 TMPRSS2-ERG protein, we first searched our IP-MS/MS data for the modified peptides with variable N-terminal acetylation, N-terminal methionine truncation, and phosphorylation of threonine, serine and tyrosine. Our search identified peptide N-acetyl-TASSSSDYGQTSK with the near-complete series of y- and b-fragment ions (**Figure 3C**). Additional search of our IP-MS/MS data against the custom database with the unreviewed TrEMBL sequences confirmed the absence of any other non-standard isoforms. We then confirmed the N-terminal identity of the T1/E4 TMPRSS2-ERG protein with more sensitive IP-SRM assays using six different synthetic peptide internal standards (**Figure 3A)**, including peptides with phosphorylated threonine or serine, as previously suggested for ERG over-expressed in 293T cells.^35^ IP-SRM confirmed the presence of N-acetyl-TASSSSDYGQTSK peptide, but no phosphorylation of threonine or serine (**Figure 3B**).

**Figure 3.**
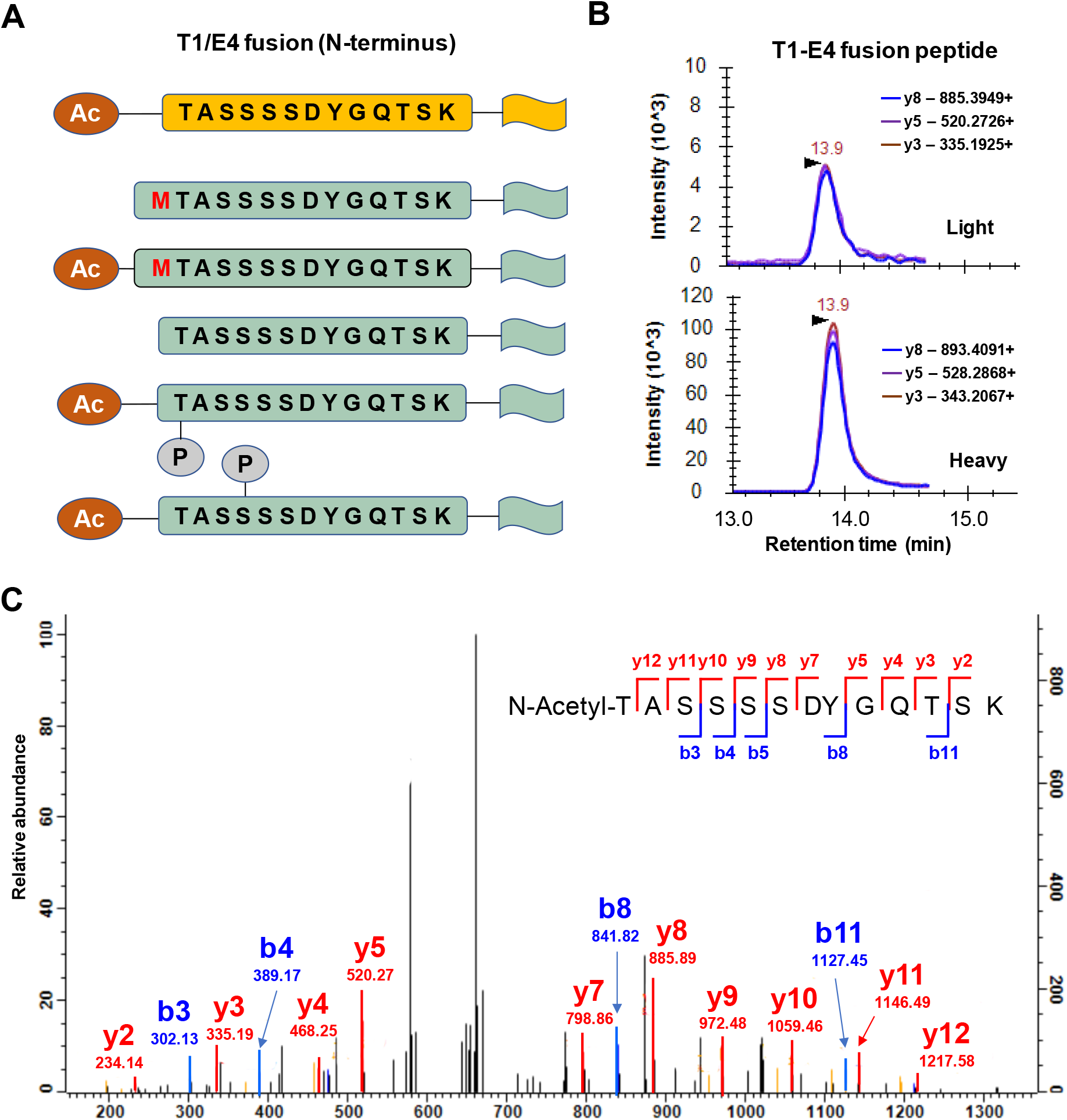
Detection of the unique fusion peptide of TMPRSS2-ERG protein in VCaP cells. **(A)** Six different N-terminal peptides, including N-term acetylated (Ac) and phosphorylated (P) peptides, were measured by SRM. **(B)** IP-SRM measurement of the unique fusion peptide N-acetyl-TASSSSDYGQTSK in the VCaP lysate. This peptide represented two most abundant isoforms T1E4-ERG and T1E4-ERG_Δ7b (∼85% total ERG). **(C)** MS/MS spectrum of N-acetyl-TASSSSDYGQTSK peptide identified in the VCaP lysate by shotgun IP-MS.

### Identification of ERG Interactome

We hypothesized that orthogonal IP-MS/MS assays with the N-term and C-term antibodies could facilitate elucidation of the exhaustive interactome of endogenous ERG (**Figure 1B**). We first tested several previously proposed mild and nondenaturing detergents,^24^ including (i) 0.1% Rapigest; (ii) modified IP lysis buffer (0.1% Rapigest, 1% NP-40, 0.1% sodium deoxycholate); and (iii) 0.5% SDS mixed with 0.5% NP-40. As a result, we selected 0.1% Rapigest, based on the completeness of the cell lysis, total protein amount, and ERG recovery.

Following simultaneous immunoprecipitation on the same 96-well plate and proteomic sample preparation, ERG interactomes were identified by shotgun MS with the label-free quantification. MaxQuant search against the “reviewed” UniProtKB human protein database resulted in identification and quantification of 449 proteins (**Table S9**). Proteins identified with false detection rate (FDR) of 1.0% and with fold change >1.3 and P-value <0.05 were selected as ERG-interacting proteins (**Table S10**). As a result, IP-MS with the C-term antibody identified 29 ERG-interacting proteins (**Table S10**), including the BAF (SWI/SNF) chromatin remodeling complex subunits, androgen receptor,^36^ and some nuclear receptor co-activators including NCOA2_HUMAN and NCOA6_HUMAN (**Figure 4A)**. Identification of eight proteins of the BAF complex (∼2,000 kDa polymorphic assemblies of ∼14 subunits encoded by 28 genes^37^) with significant fold changes (P-value <0.05) was in good correlation with a recent study by Sandoval et al.^24^ Our interactome also included the Ewing’s Sarcoma breakpoint protein EWS_HUMAN (**Figure 4A**), which was reported to specifically interact with ERG and other oncogenic ETS factors, promote gene expression, and drive oncogenic transformation and cell migration of prostate cells.^38^ We believe that in our study we identified one of the most comprehensive ERG interactomes (**Figure 4D and Figure S5**).

**Figure 4.**
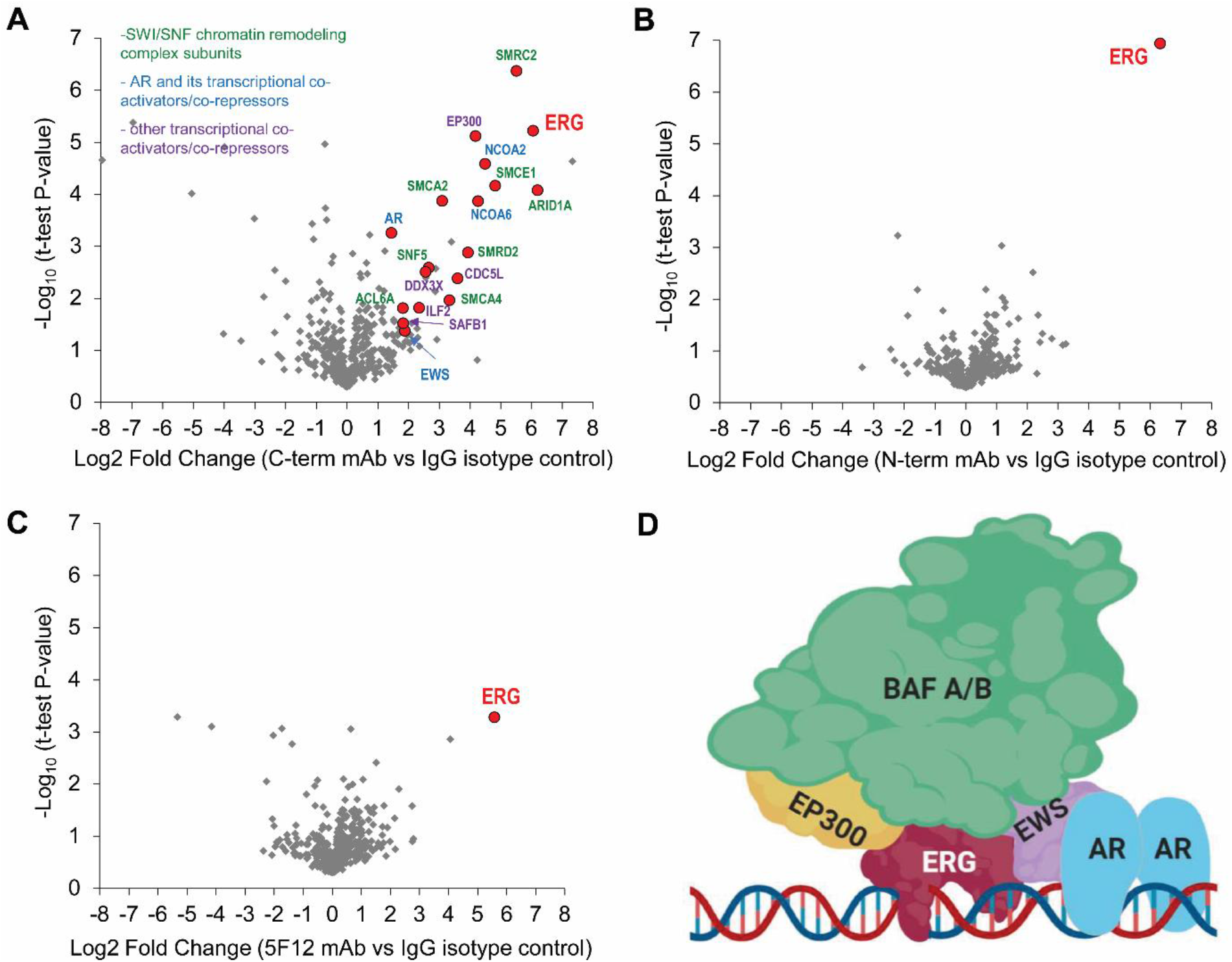
TMPRSS2-ERG interactome identified in VCaP cells. IP experiments were completed with the C-term mAb **(A)**, N-term mAb **(B)** and 5F12 mAb **(C)**. IP was completed simultaneously on a 96-well plate, and IP conditions, proteomics sample preparation and the VCaP lysate were identical. Volcano plots presents proteins identified by IP-MS. **(D)** Schematic presentation of some ERG-interacting proteins, including BAF complex proteins (green), in agreement with our study and literature data.

Interestingly, while ERG protein alone has been efficiently enriched 81-fold (N-term mAb, **Table S11**), 67-fold (C-term mAb), and 48-fold (5F12 mAb), immunoprecipitation with the N-term mAb and 5F12 mAb (unknown but likely an internal epitope) failed to identify any ERG-interacting proteins (**Figure 4B and 4C)**. Such unexpected result suggested that some antibodies could interfere with or disrupt protein-protein interactions of ERG. Similar case on antibody-mediated disruption of protein-protein interactions we have previously observed for the TEX101-DPEP3 complex.^39^ This unexpected finding led us to the hypothesis that the ERG interactome could be disrupted by some short synthetic peptides representing the known 9FY epitope, thus paving the road to development of targeted anti-ERG therapies.

### Independent verification of ERG-interacting proteins by PRM

To verify some ERG-interacting proteins, we developed parallel reaction monitoring (PRM) assays for the most interesting ERG-interacting proteins, including some BAF complex proteins (ARI1A, SMCA2 and SMCE1), androgen receptor (ANDR), nuclear receptor co-activators (NCOA2 and NOCA6), and a transcriptional co-regulator EWS (EWSR1 gene). Targeted assays with the heavy peptide internal standards facilitated precise measurements of protein relative abundances and more accurate estimation of the enrichment fold-changes, as previously demonstrated.^25, 40^ Interestingly, independent verification revealed potentially three groups of ERG-interacting proteins: 1) strong and moderate ERG binders disrupted by the N-term antibody (NCOA2, NCOA6 and ARI1A); 2) a strong ERG binder not disrupted by the N-term antibody (EWS); and 3) weak ERG binders or potential false-positives (**Table S11**).

### Evaluation of ERG interactome disruption by the N-term epitope peptides

Since the N-term antibody could interfere with the ERG interactome, we hypothesized that synthetic peptides which represented the minimal epitope (RVPQQDWL) or an extended sequence (KMSP<u>RVPQQDWL</u>SQ)^41^ could specifically compete and disrupt the interaction between ERG and its interactome.

First, we confirmed by IP-PRM that both synthetic peptides had no effect on the ERG capture by the C-term antibody, but disrupted the ERG capture by the N-term antibody to the similar extent. Second, we selected only the minimal epitope peptide RVPQQDWL, and evaluated its impact on the ERG-interacting proteins. As a result, only some weak effects were measured at a very high 800 µM concentration of RVPQQDWL (**Figure 5A, Table S12**). Since such concentration would account for the EC50 in the mM range, if any at all, we concluded that the minimal epitope peptide was not a candidate for the disruption of ERG-interacting proteins.

**Figure 5.**
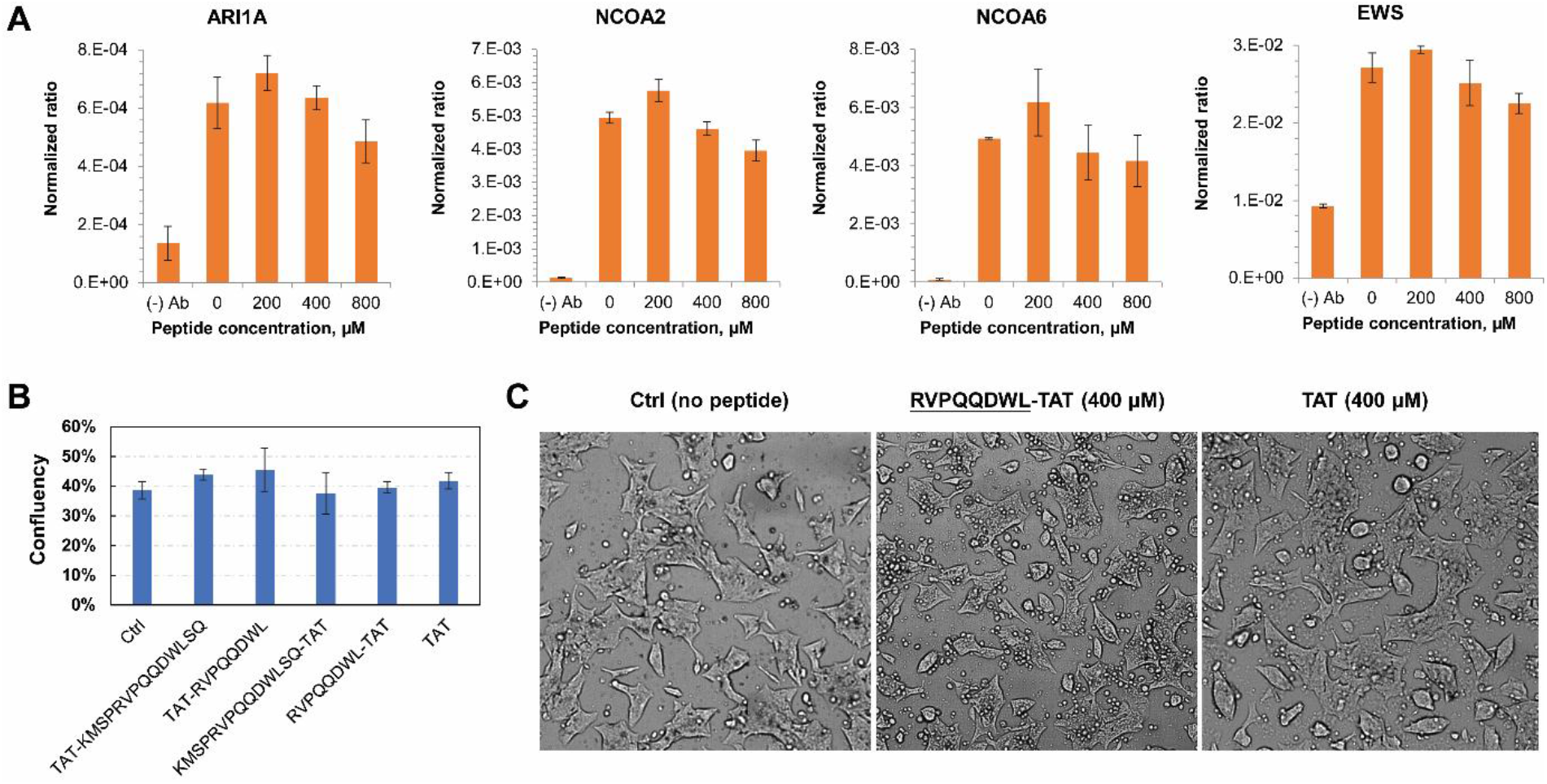
Assessment of the ERG interactome disruption by synthetic 9FY epitope peptides. **(A)** ERG-interacting proteins were enriched from the VCaP lysate using the C-term mAb, incubated with the increasing amounts of 9FY epitope peptide RVPQQDWL, and quantified by PRM assay. No significant differences were observed. **(B)** Confluency of VCaP cells treated with 400 μM epitope peptides (including peptides with the cell-permeable HIV-TAT sequences) and grown for 4 days. **(C)** Comparison of VCaP cell morphology untreated, treated with 400 μM RVPQQDWLSQ-TAT and TAT peptides for 4 days. No difference in cell morphology was observed.

Interestingly, our data suggested that some ERG-interacting proteins (NCOA2, NCOA6 and the BAF complex proteins) were binding ERG somewhere between the N-term epitope and the ETS DNA-binding domain (251 aa region), most probably near the 88 aa PNT domain located in that region (**Figure S5**). Disruption of the ERG interactome in this region was probably due to the steric hindrance between the large 160 kDa antibody and very large transcriptional regulators NCOA2, NCOA6 and EP300 (160-260 kDa), or extremely large BAF complexes (∼2,000 kDa). On the contrary, interaction of a relatively small EWS protein (68 kDa) binding ERG in the region between the ETS domain and the C-terminal epitope,^38^ was barely affected by the N-term antibody in our verification phase **(Table S11)**.

### Impact of the N-term epitope peptides on proliferation and morphology of VCaP cells

In parallel with evaluation of the N-term epitope peptide impact by PRM assays, we evaluated the impact of these peptides on proliferation and morphology of VCaP cells. The cell-permeable N-term epitope peptides were conjugated to the cationic HIV-TAT motif GRKKRRQRRRG to facilitate uptake into VCaP cells, as previously reported.^42^ We found that 400 µM of the cell-permeable N-term epitope peptides had no impact on VCaP proliferation (**Figure 5B, Table S13**) or on VCaP morphology (**Figure 5C**).

## Discussion

Proteomics by mass spectrometry has advanced to the level of identification and quantification of nearly whole proteomes of human cells (∼12,000 proteins per cell).^43^ Without additional approaches for protein fractionation or enrichment, shotgun proteomic methods are still limited in terms of quantification of low-abundance cellular proteins, such as transcription factors or cell signaling receptors.

IP-MS and IP-SRM assays have recently gained considerable interest due to their high sensitivity and selectivity for quantification of low-abundance proteins in cells and biological fluids.^44^ Immunoprecipitation substantially reduces sample complexity and facilitates quantification of multiple peptides per protein, thus enabling in-depth analysis of post-translational modifications and resolution of splicing isoforms. In our previous studies, IP-SRM assays have been utilized to quantify low-abundance kallikrein-related peptidases,^45^ resolve protein isoforms,^46^ screen for antibody clones,^47^ discover the TEX101-DPEP3 protein complex,^39^ and detect a low-abundance missense variant of TEX101 protein.^48^ In this study, we focused on development of IP-MS and IP-SRM assays for the low-abundance TMPRSS2-ERG fusion protein, which triggers malignant transformation of prostate epithelial cells in nearly 50% of prostate cancer cases.^49^ TMPRSS2-ERG mRNA expression has been well characterized, but its association with the more aggressive cancer^14, 15^ is still conflicting. Here, we hypothesized that IP-MS and IP-SRM could facilitate quantification of the low-abundance TMPRSS2-ERG protein, resolve the identity and abundance of TMPRSS2-ERG isoforms, elucidate N-terminal post-translational modifications, and discover the ERG interactome. We thought that the development of novel assays and detailed characterization of TMPRSS2-ERG fusion at the protein level could enable future translational and pre-clinical studies on its association with the aggressive prostate cancer.

Wild-type ERG is a potent transcription factor and is expressed at low-to-nothing levels in healthy tissues, normal human prostate stromal cells (PrSCs) and the majority of prostate cancer cell lines. Oncogenic nature of ERG protein is exerted through its function as a transcription factor which promotes cell migration and cancer progression.^50^ Only two prostate cancer cell lines harboring TMPRSS2-ERG gene fusion express measurable amounts of ERG transcripts: VCaP (∼50,000-fold higher than baseline expression in PrSCs) and NCI-H660 (∼6,000-fold higher than baseline expression).^51^ The most powerful mass spectrometry instruments are now able to detect multiple tryptic peptides of ERG protein, but still cannot fully resolve ERG isoforms.^43^ He et al. have previously pioneered quantification of ERG protein by combining two-dimensional chromatography with SRM assays, but did not quantify relative abundances of splicing isoforms or detected the unique TMPRSS2-ERG fusion peptide.^29, 30^ The level of ERG in VCaP cells has been estimated at 1.8 fg per cell, which was in agreement with our IP-SRM data (2.2 fg, or ∼27,000 copies per cell).

It has previously been demonstrated that distinct protein splicing isoforms may have unique molecular functions and even exhibit dominant-negative effects,^52^ such as the N-terminally truncated isoform ERα46 of the estrogen receptor alpha.^53^ The identity and abundance of splicing mRNA isoforms could now be readily measured by RNA-sequencing in cell lines^31^ and human tissues.^54^ Elucidation of the identity, abundance and function of the distinct protein isoforms is still challenging and could be considered as a later milestone for the bottom-up and top-down proteomics, as well as proteogenomics.^55-57^

ERG mRNA isoforms were previously well characterized in cells and prostate cancer tissues.^58^ Interestingly, the shortest mRNA isoform-8 encoding T1E4-ERG_7bpA protein lacking DNA-binding ETS domain has been suggested to exhibit dominant-negative effects on the ERG-mediated gene expression. Interestingly, higher abundance of isoform-8 mRNA versus total ERG has even been linked to the more favorable pathology and outcomes of prostate cancer.^12^ This isoform could compete for ERG homodimerization or protein-protein interactions, but the resulting complex would fail to trigger transcriptional activation.^19, 20^ In our present study, IP-SRM assays resolved for the first time the relative abundance of four ERG isoforms in VCaP cells, and unambiguously demonstrated that isoform-8 was expressed at mRNA, but not at the protein level (**Figures S6 and S3**).

Recently, it has been demonstrated that degradation of the wildtype ERG protein is mediated through the SPOP binding site 42-AS<u>S</u>S<u>S</u>-46 (“degron”) located at exon 4. Truncated T1E4-ERG isoform displayed a significantly reduced interaction with SPOP ubiquitin ligase *in vivo* and *in vitro*, and Ser44 or Ser46 were found as major sites for CKI-dependent phosphorylation.^33, 35^ The N-term truncation of ERG was proposed as the major mechanism leading to the conformational change and reduced binding of SPOP to the ERG degron. The fact that the N-terminal post-translational modifications could define ERG stability^34, 59^ encourage us to characterize the N-terminal composition of the endogenous TMPRSS2-ERG fusin protein in VCaP cells. Our SRM measurements of the N-term peptide N-acetyl-<u>T</u>AS<u>S</u>SSDYGQTSK revealed that Thr1 and Ser4 were not phosphorylated. Our study revealed that the two most abundant isoforms T1E4-ERG and T1E4-ERG_Δ7b (85% of total ERG) were also modified by the N-term methionine truncation and threonine acetylation, which could further reduce SPOP binding and contribute to the prolonged ERG protein half-life in VCaP cells. Finally, Uniprot search revealed that none of the human protein isoforms had the N-term motif MTASSS, one isoform had the motif MTASS (Q9H3U1-2; minor expression in testis tissues), and only five proteins had the N-term motif MTAS (Q9NV35, Q9H596, Q6NXG1, P31271, Q9NQ50). Development of affinity reagents specifically recognizing the N-terminal protein sequence N-acetyl-TASSSS could become a strategy to target TMPRSS2-ERG fusion protein for degradation,^60^ thus enabling the next generation of targeted therapies of prostate cancer.

In addition, we applied our orthogonal IP-MS assays to identify TMPRSS2-ERG interactome. It should be noted that the transcriptional activation by ERG could be mediated through its dimerisation^19^ and complex interplay with other transcriptional factors and regulators.^22, 38^ For example, ERG interaction with androgen receptor has been previously identified by co-immunoprecipitation-western blotting.^22, 61^ Two previous IP-MS studies completed with potentially lower-confidence polyclonal anti-ERG antibodies identified probably only partial ERG interactomes (for example, lacking EWS^24^ or majority of BAF complex proteins^62^). Interestingly, co-IP and western blotting with ERG antibodies specific for distinct epitopes^24^ revealed significant enrichment of SMARCA4 (BRG1 gene) with the C-terminal polyclonal antibody, but very low amounts of SMARCA4 enrichment with the monoclonal antibody targeting the internal region of ERG, thus suggesting that antibodies targeting internal regions of ERG could interfere with its interactome. We believe that our IP-MS assay with the C-term monoclonal antibody identified the most comprehensive ERG interactome (including BAF complexes, transcriptional regulators, EWS and other proteins).

We have previously demonstrated that some high-affinity monoclonal antibodies could disrupt protein-protein interactions.^39^ Likewise, our present data suggested that the N-term monoclonal antibody could interfere with binding of some ERG-interacting proteins, and that some ERG-interacting proteins (especially very large ∼2,000 kDa BAF complexes and 150-260 kDa transcriptional regulators) were binding to the 251 aa region between the N-term epitope and ETS DNA-binding domain. These large proteins were probably displaced by the N-term epitope antibody and the full ERG antibody, but not by the C-term antibody. On the contrary, a relatively small 68 kDa EWS protein binding within the region between ETS domain and the C-term epitope was barely affected. In future, exact regions of protein-protein interactions could be determined using series of recombinant truncated ERG proteins, epitope mapping arrays, or cross-linking mass spectrometry.^63^ It would be extremely interesting to investigate if a shorter and quite abundant isoform T1E4-ERG_Δ7b (lacks 24 aa sequence between the N-term epitope and ETS domain; 14% of total ERG) could have differential binding for partner proteins crucial for the ERG transcriptional activity. This isoform could thus have dominant-negative effects through the protein-protein interaction domain, similar to the hypothetical dominant-negative activity through the ETS domain proposed for the undetected T1E4-ERG_7bpA isoform (isoform-8 mRNA). In future, our IP-SRM assays could also be utilized to quantify distinct ERG isoforms and their interactomes in clinical samples, including tissues, urine, seminal plasma or exfoliated prostate epithelial cells, thus contributing to the top research need on precision diagnostics prioritized by the Movember Prostate Cancer Landscape Analysis.^64^ Finally, ERG and some BAF complex proteins are now being investigated as therapeutic targets in prostate cancer.^65^ Short peptides efficiently disrupting ERG protein-protein interactions, similar to peptides disrupting ERG-DNA interactions^42^, could be identified and emerge as the next generation targeted therapies of prostate cancer.

## Conclusions

To conclude, in this study we developed IP-SRM and IP-MS assays for the quantification of a low abundant transcriptional factor TMPRSS2-ERG fusion protein, its isoforms, and its interactome in VCaP cells. Our orthogonal IP-SRM assays quantified for the first time the relative abundance of four isoforms, and revealed that the T1E4-ERG isoform accounted for 71% of the total TMPRSS2-ERG protein in VCaP cells. For the first time, the N-terminal peptide (methionine-truncated and N-acetylated TASSSSDYGQTSK) unique for the T1/E4 fusion was identified. The interactome data suggested that TMPRSS2-ERG protein-protein interactions, including SWI/SNF chromatin remodeling complex subunits and numerous transcriptional co-regulators, were exerted through at least two different regions. Our sensitive and selective IP-SRM assays present alternative tools to quantify TMPRSS2-ERG fusion protein, its isoforms and interactomes in clinical samples, thus paving the road to development of more accurate biomarkers and diagnostics of prostate cancer. The precise knowledge on the molecular identity and interactomes of the endogenous TMPRSS2-ERG fusion protein facilitates development of novel targeted therapies of prostate cancer.

## Experimental procedures

### Hypothesis, study design and objectives

We hypothesized that the use of N-term and C-term anti-ERG antibodies with IP-MS assays will provide novel knowledge on the identity and abundance of ERG isoforms and interactome. Our study was designed to measure isoforms and interactome of the endogenous TMPRSS2-ERG fusion protein in VCaP cells with novel IP-MS and IP-SRM assays. Orthogonal N-term and C-term monoclonal anti-ERG antibodies were employed for the enrichment of ERG protein, its isoforms and ERG-interacting proteins. Specific objectives included: (1) develop novel IP-MS and IP-SRM assays; (2) quantify the relative abundance of TMPRSS2-ERG protein isoforms, (3) identify the N-term modifications of TMPRSS2-ERG protein, (4) identify TMPRSS2-ERG protein interactome in VCaP cells, and (5) investigate if synthetic N-term epitope peptides can disrupt TMPRSS2-ERG interactome.

### Chemicals and Cell Culture Reagents

Iodoacetamide, dithiothreitol, L-methionine and trifluoracetic acid (TFA) were purchased from Thermo Fisher Scientific (Burlington, ON, Canada). Optima water and acetonitrile (ACN) were purchased from Fisher Scientific (Fair Lawn, NJ). Formic acid (FA) was obtained from Sigma-Aldrich (Oakville, ON). Stable isotope labeled peptides (SpikeTides_L and SpikeTide_TQL) were obtained from JPT Peptide Technologies GmbH (Germany). VCaP and LNCaP prostate cancer lines were obtained from the American Type Culture Collection (Burlington, ON, Canada). Cell lines were cultured in a humidified incubator at 37 °C and 5% CO_2_. Dulbecco’s Modified Eagles Medium (HyClone) and RPMI 1640 medium (Gibco) were used to culture VCaP and LNCaP cells, respectively. Mediums were supplemented with 10% fetal bovine serum (Invitrogen) and 1% penicillin-streptomycin (Invitrogen) prior to use.

### Cell Lysis and Sample Preparation

Cell pellets were lysed in 50 μL 0.1% RapiGest SF (Waters, Milford, MA) with repeated pipetting, vortexing and probe sonication at 20 kHz. EDTA-free protease inhibitor cocktail (Roche) combined with Benzonase™ Nuclease (Fisher Scientific) were added prior to cell lysis to reduce proteolysis and digest nucleic acids. To remove cell debris, lysates were centrifuged at 16,000 g and 4 °C for 10 min. Total protein of lysates was measured by Pierce BCA protein assay kit (Fisher Scientific). Proteins were denaturated, and disulfide bonds were reduced by 10 mM dithiothreitol at 70 °C for 15 min, and alkylated with 20 mM iodoacetamide at RT in the dark for 45 min. Digestion was performed overnight at 37 °C using recombinant dimethylated SOLu-trypsin (Sigma-Aldrich) with a trypsin:protein ratio 1:20. TFA (1%) was added to cleave and precipitate Rapigest SF, and 1 μL of 0.4 M L-methionine was added to the peptide digest, to limit methionine oxidation during storage. Prior to liquid chromatography, 10 μL OMIX C18 tips (Agilent Technologies) were used for desalting and microextraction of tryptic peptides. Finally, samples were diluted in 5% ACN with 0.1% FA.

### RT-PCR

Total RNA was extracted from LNCaP and VCaP cells using TRIzoL (ThermoFisher Scientific). RNA was reversely transcribed to cDNA via iScriptTM Reverse Transcription Supermix (Biorad). After quantification by NanoVue Plus spectrophotometer (GE Healthcare), 500 ng cDNA was used as PCR template for the amplification of TMPRSS2-ERG fusion. Hot Start Taq 2X Master Mix (New England Biolabs) and GeneAmp PCR System 2700 thermal cycler (Applied Biosystems) were used. The forward and reverse PCR primers for T1/E4 fusion were 5’-TAGGCGCGAGCTAAGCAGGAG-3’ and 5’-CCATAT TCTTTCACCGCCCACTCC-3’ (Integrated DNA Technologies), respectively. The primer pair for ERG isoform 8 was 5’-GGTACGAAAACACCCCTGTG-3’ (forward) and 5’-CCAAATCAACAGAGGCAGAA-3’ (reverse), and those for total ERG was 5’-AACGAGCGCAGAGTTATCGT-3’ (forward) and 5’-GTGAGCCTCTGGAAGTCGTC-3’ (reverse). The final volume was 25 μL, and an initial denaturation step of 95°C for 5 min was followed by 40 cycles of 30 s at 95 °C, 30 s at 59 °C, 30 s at 72 °C, and one cycle at 72 °C for 5 min. Formation of the cDNA products of T1/E4 fusion gene (**Supplemental Figure S1**), ERG isoform 8 and total ERG was confirmed by 2% agarose gel electrophoresis.

### Immunoprecipitation

A rabbit monoclonal anti-ERG antibody EPR3864(2) (C-term epitope PNTRLPTSHMPSH; Abcam, Toronto, ON), a mouse monoclonal antibody 9FY (N-term epitope KMSPRVPQQDWLSQ; BioCare Medical, Markham, ON) and a mouse monoclonal antibody OTI5F12 (likely an internal epitope since the full 479 aa sequence was used as an antigen; Origene Technologies, Rockville, MD) were used as capture antibodies for the enrichment of ERG protein from cell lysates (**Figure 1B**). Two μL of antibody diluted in PBS (pH 7.4) was coated on a high-binding 96-well polystyrene microplate (Greiner Bio-One) with 100 μL per well and incubated overnight at room temperature (RT). As 9FY antibody was provided in Renoir Red solution with carrier proteins, a goat anti-mouse Fcγ fragment specific antibody (Jackson Immunoresearch Labs) was first coated on the plate for 9FY pull-down. After washing 3 times (200 μL each) with the wash buffer (0.1% Tween 20 in PBS), the plate was blocked for 1 h at RT by adding 200 μL blocking buffer (2% BSA in wash buffer) into each well. The washing step was repeated, followed by addition of 80 μg total protein VCaP lysates into each well and diluting to 100 μL with dilution buffer (0.1% BSA in wash buffer, 0.2 μm filtered). After 2 h incubation with continuous shaking, the plate was finally washed 3 times with wash buffer and 3 times with 50 mM NaHCO_3_. Trypsin (0.25 ng per well) was used for digestion, as previously described. The heavy isotope-labeled peptide standards were mixed and diluted to the final concentration of 100 fmol/μL. Three μL of the internal standard mixture were spiked to each sample before (SpikeTides_TQL peptides) or after (SpikeTides_L peptides) trypsin digestion and each digest was analyzed in triplicates. For the interactome studies, antibody isotype controls included (i) anti-FOLH1 mouse monoclonal antibody (clone 3B5, Abnova) as an IgG_1_ isotype control for OTI5F12 and 9FY antibodies; (ii) anti-KLK3 rabbit monoclonal antibody (Sino Biological) as an IgG isotype control for EPR3864(2) antibody.

### Selection of Tryptic Peptides and Development of SRM Assays

Peptide Atlas, neXtProt database and in-house IP-LC-MS/MS data were used to select best peptides for the total ERG, or peptides unique and shared by specific isoforms. Excision of the N-term methionine, N-term acetylation, and serine or threonine phosphorylation were considered for the unique N-term peptide MTASSSSDYGQTSK of the T1/E4 fusion. In total, 14 synthetic heavy isotope labeled SpikeTides_L peptides were used as internal standards for SRM assay development (**Table S2**). Initially 28 heavy and light peptide pairs (280 transitions) were included into a multiplex unscheduled SRM method with 5 ms scan times per transition. Retention times and heavy to light ratios of the transitions were monitored, and optimization of transitions was performed by removing transitions with low selectivity and possible interferences in the cell lysate. Redundant poorly-performing peptides were removed, and 6 peptides with 3 or 4 most intense and reproducible transitions per peptide (42 transitions) were finally scheduled within 2 min intervals in a single multiplex scheduled SRM assay. The scan time of 10 ms ensured acquisition of at least 20 points per peak. Superposition of light and heavy peptide peaks and the order of y-ion transition intensities ensured the correct identities of peptides.^66, 67^ After preliminary experiments with SpikeTides_L peptides, purified and heavy isotope labelled SpikeTides_TQL peptides with JPT-tags [serine-alanine-(3-nitro)tyrosine-glycine] were used as internal standards to facilitate absolute quantification.

### Chromatography and Targeted Mass Spectrometry

A quadrupole ion-trap mass spectrometer (AB SCIEX QTRAP 5500) coupled to EASY-nLC II (Thermo Scientific) via a NanoSpray III ion source (AB SCIEX) was used for SRM assays. The tryptic peptides were loaded at 5 μL/min onto a C18 trap column (Thermo Scientific, 100 µm ID×2 cm, 5 μm, 120 Å). Peptides were separated with PicoFrit columns (New Objective, 15 cm×75 μm ID, 8 μm tip, PepMap C18, 3 μm, 100 Å) and 28 min gradients (300 nL/min). The gradient started with 5% buffer B and ramped to 65% buffer B over 20 min, followed by an increase to 100% buffer B within 1 min, and continued for 7 min. QTRAP 5500 parameters were: 2300 V ionspray; 75 °C source temperature; 2.0 arbitrary units for gas 1 (N_2_), 0 arbitrary units for gas 2; 25 arbitrary units for curtain gas (N_2_); and 100 V declustering potential. Q Exactive Hybrid Quadrupole-Orbitrap (Thermo Scientific) coupled to EASY-nLC 1000 (Thermo Scientific) was used for PRM assays. The mobile phase consisted of 0.1% FA in water (buffer A) and 0.1% FA in ACN (buffer B). Acclaim PepMap 100 nanoViper C18 column (Thermo Scientific, 100 µm ID×2 cm, 5 μm, 100 Å) was used as pre-column for sample loading, while the EASY-Spray C18 column (Thermo Scientific, 15 cm×75 μm ID, 3 μm, 5 μm) was used as the analytical column. An 18-min gradient (400 nL/min) started with 0% buffer B and ramped to 50% buffer B over 15 min, followed by an increase to 100% buffer B within 1 min, and kept for 2 min. PRM scans were performed at 17.5 K resolution with 27% normalized collision energy. AGC (Automatic Gain Control) target value was set to 3×10^6^ with a maximum injection time of 100 ms and an isolation width of 2.0 m/z. The performance of the nanoLC-MS systems was ensured every 6 runs using 10 μL injections of 20 fmol/μL BSA.

### Chromatography and Shotgun Mass Spectrometry

ERG interactome was identified using Orbitrap Elite™ Hybrid Ion Trap-Orbitrap mass spectrometer (Thermo Scientific) coupled to EASY-nLC II. The peptides were eluted at 300 nL/min using a 2-hour gradient: 5% B for 5 min, 5-35% B for 95 min, 35-65% B for 10 min, 65-100% B for 1 min and 100% B for 9 min. All LC-MS/MS data were acquired using XCalibur (v. 2.2). The MS1 scans (400-1250 m/z) were performed at 60 K resolution in the profile mode, followed by top 20 ion trap centroid MS/MS, acquired at 33% normalized collision energy. FTMS ion count was set to 1×10^6^ with an injection time of 200 ms, while MS/MS scans were set to 9,000 count and 100 ms injection time. MS/MS acquisition settings included 500 minimum signal threshold, 2.0 m/z isolation width, 10 ms activation time, and 60 s dynamic exclusion. Monoisotopic precursor selection and charge state rejection were enabled, +1 and unknown charge states were rejected. Instrument parameters included 230 °C capillary temperature, 2.0 kV source voltage and 67% S-lens RF level.

### IP-SRM Assessment of Interactome Disruption by Synthetic 9FY Epitope Peptides

Synthetic 9FY epitope peptides (HCl salts; Biomatik) included the minimal (RVPQQDWL) and extended (KMSPRVPQQDWLSQ) epitopes. C-term antibody EPR3864(2) (1 μg per well) was used to enrich ERG interactome from VCaP lysate (80 μg total protein). Following ERG interactome enrichment on 96-well plates (analytical duplicates), 9FY epitope peptides were added at 0, 200, 400 and 800 µM (final in 100 μL PBS) and incubated overnight at RT, followed by three time washing and trypsin digestion (0.25 ng per well). SpikeTides_L peptides (200 fmol per well) were used for the accurate relative quantification of seven ERG-interacting proteins (**Table S3**). Each digest was analyzed in technical duplicates with PRM using Q Exactive.

### MS data analysis

Raw files of SRM and PRM experiments were analyzed using Skyline Targeted Proteomics Environment v20.1.0.76 (MacCoss Lab).^68^ Peak boundaries were adjusted manually, and the integrated areas of all transitions for each peptide were extracted. Light-to-heavy peak area ratios were used for accurate relative or absolute quantification of endogenous peptides. Shogun MS data were search against a nonredundant reviewed human UniProtKB/Swiss-prot database (20,365 entries) using MaxQuant software (v1.6.7.0).^69^ Search parameters included: trypsin enzyme specificity, 2 missed cleavages, 7 aa minimum peptide length, top 8 MS/MS peaks per 100 Da, 20 ppm MS1 and 0.5 Da MS/MS tolerance. Variable modifications included methionine oxidation, N-terminal acetylation and deamidation (N). False-discovery rate (FDR) was set to 1% at both protein and peptide levels. Label-free quantification (LFQ) algorithm was used for quantification. MaxQuant proteinGroups.txt file was uploaded to Perseus software (version 1.6.12.0)^70^ for further analysis. Protein identifications marked as “Only identified by site”, “Reverse” and “Contaminants” were excluded. LFQ intensities were log2-transformed and missing LFQ values were imputed with values representing the normal distribution (0.2 width and 1.8 down shift). Log2 fold change of 1.3 and one-tail t-test P-value<0.05 was applied to determine proteins statistically enriched by anti-ERG antibodies versus isotype controls. In addition, IP-shotgun MS data was searched against the custom database with all possible ERG isoforms present in the unreviewed TrEMBL database (accessed on Mar. 30, 2020). Raw shotgun MS data have been deposited to the ProteomeXchange (PXD021236; username; password: mpNlP18V), while raw SRM and PRM data have been deposited to Peptide Atlas with the dataset identifier PASS01624 (www.peptideatlas.org/PASS/PASS01624 or ftp://PASS01624:SL727hk@ftp.peptideatlas.org).

### Assessment of Synthetic 9FY Epitope Peptides in Live VCaP Cells

To facilitate cell penetration and investigate synthetic 9FY epitope peptides in live VCaP cells, RVPQQDWL and KMSPRVPQQDWLSQ peptides were synthesized with the TAT sequence (GRKKRRQRRRG) at N- and C-terms (HCl salts; Biomatik). Peptides (final 400 μM in PBS) were incubated with VCaP cells (15,000 cells per well) on a 96-well plate. After 4 days incubation, the cell morphology was monitored via live cell imaging using the MetaXpress XLS system (Molecular Devices). Then the relative confluency of cells in each well was estimated based on the averaged cell counting of four images. The cell counting was performed using the cell segmentation algorithm in ImageJ (FIJI) software (version 1.51w).^71^

## Acknowledgements

We thank Prof. Xingfang Li for the access to QTRAP 5500 mass spectrometer and Dr. Konstantin Stoletov for providing the VCaP and LNCaP cells.

## Author contributions

A.P.D. designed the research project. Z.F. performed all major experiments, analyzed data and wrote manuscript. Y.R. assisted with cell culture and sample preparation. A.P.D., Z.F. and C.X.L. edited manuscript.

## Supplemental materials

This article contains supplemental materials.

## Non-standard abbreviations

aa: amino acids
CV: coefficient of variation
FDR: False discovery rate
FWHM: Full width at half maximum
ELISA: Enzyme-linked immunosorbent assay
LC-MS/MS: liquid chromatography - tandem mass spectrometry
LFQ: Label-free quantification
PCa: Prostate cancer
PRM: parallel reaction monitoring
PSA: Prostate-specific antigen
SRM: Selected reaction monitoring
ERG: v-Ets avian erythroblastosis virus E26 oncogene homolog

## Table of Contents Figure

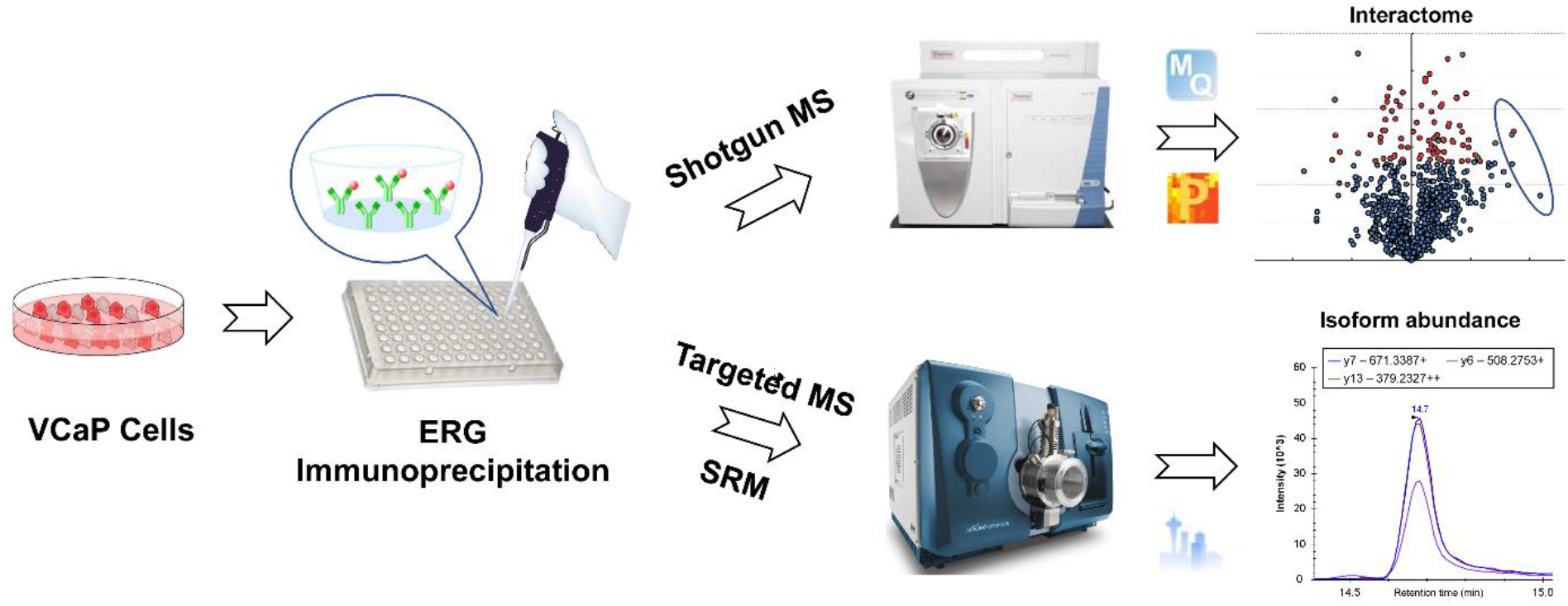

