## Supplemental Text and Figures for "Mapping Isoform Abundance and Interactome of the Endogenous TMPRSS2-ERG Fusion Protein with Orthogonal Immunoprecipitation-Mass Spectrometry Assays"

#### Calculation of the relative abundance of T1-E4 TMPRSS2-ERG isoforms in VCaP cells

*Assumptions:*

1) Using quantifiable internal standard peptides with the trypsin-cleavable tags, we can accurately estimate "absolute" molar amounts of the endogenous ERG peptides.

2) Without IP, we could not detect by SRM the mutually exclusive peptides ITTRPDLPYEPPR and NTDLPYEPPR. Furthermore, IP with the C-tem mAb alone could not unambiguously resolve isoform abundances. It was only IP with the N-term mAb and ITTRPDLPYEPPR / NTDLPYEPPR ratio which enabled unambiguous quantification of individual ERG isoforms.

3) We observed substantially lower, but consistent, molar amounts of ITTRPDLPYEPPR and NTDLPYEPPR peptides in all experiments, assuming potentially lower efficiency of digestion of the endogenous ERG at this position (possible missed cleavages due to additional R residues, e.g. ITTRPDLPYEPPR.R and NTDLPYEPPR.R). However, since the C-terminal sequences of both peptides were identical, we assumed that the efficiency of digestion at this position was exactly same, and used in our calculations the molar ratio of these peptides instead of the molar sum, both for the C-term and N-term data.

| **Shared and unique peptides** | **ERG isoforms** | **C-term mAb** *f*moles | **N-term mAb** *f*moles |
|---|---|---|---|
| VIVPADPTLWSTDHVR | Total ERG (all 5 isoforms) | **0.0125** | 0.0153 |
| VPQQDWLSQPPAR | T1E4-ERG (a); T1E4-ERG_Δ7b (b); T1E4-ERG_7bpA (e) | **0.0106** | 0.0194 |
| ITTRPDLPYEPPR | T1E4-ERG (a); T1E4-ERG_Δ4 (c) | **0.0039** | **0.0035** |
| NTDLPYEPPR | T1E4-ERG_Δ7b (b); T1E4-ERG_Δ4Δ7b (d) | **0.0007** | **0.0007** |
| TPLCDLFIER | T1E4-ERG_7bpA (e) | 0 | **0** |

*Data for the C-term mAb:*
1) a+b+c+d+e=0.0125 fmoles
2) e=0, thus a+b=0.0106 fmoles
3) c+d=0.0125 - 0.0106 = 0.0019 fmoles
4) (a+c)/(b+d)= 0.0039/0.0007=5.57

*Data for the N-term mAb:*
5) a/b= 0.0035/0.0007=5.0

*Calculations:*
1) a+b=0.0106 and a/b=5.0. Thus, a=0.00883 and b=0.00177 fmoles
2) c+d=0.0019 and (0.00883+c)/(0.00177+d)=5.57. Thus, c=0.00177 and d=0.00013 fmoles
3) Relative abundances are:
a [T1E4-ERG]= 71%
b [T1E4-ERG_Δ7b]= 14%
c [T1E4-ERG_Δ4]= 14%
d [T1E4-ERG_Δ4Δ7b] = 1%
e [T1E4-ERG_7bpA]= 0%

### Supplemental Figure

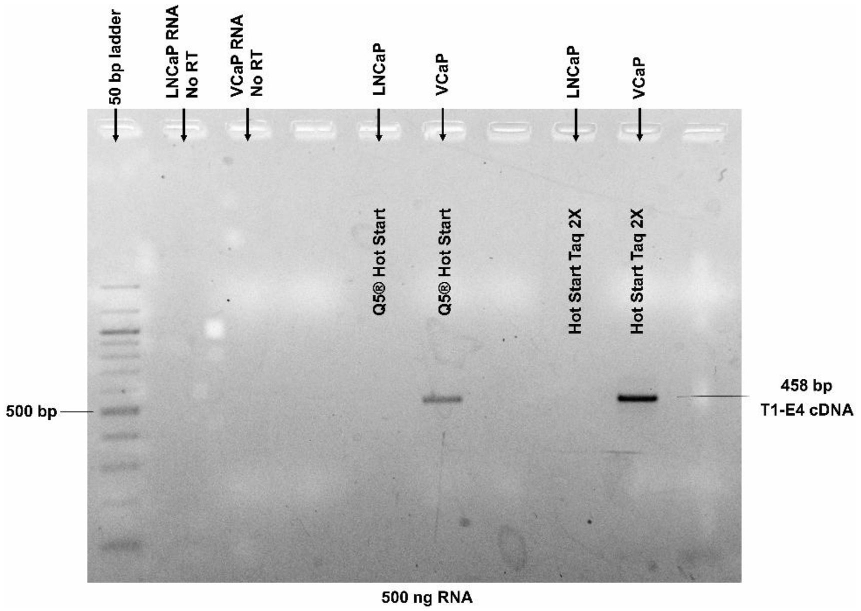


**Figure S1. Measurement of T1E4 TMPRSS2-ERG fusion mRNA in VCaP and LNCaP prostate cancer cells.** Total mRNA was extracted from VCaP and LNCaP cell lysate and reverse transcribed into cDNA, which was detected by agarose gel electrophoresis. 500 ng RNA were used as template for PCR amplification. Two PCR Hot Start Mastermix reagents (Q5® Hot Start and Hot Start Taq2X) were compared, with Taq2X polymerase having better performance.

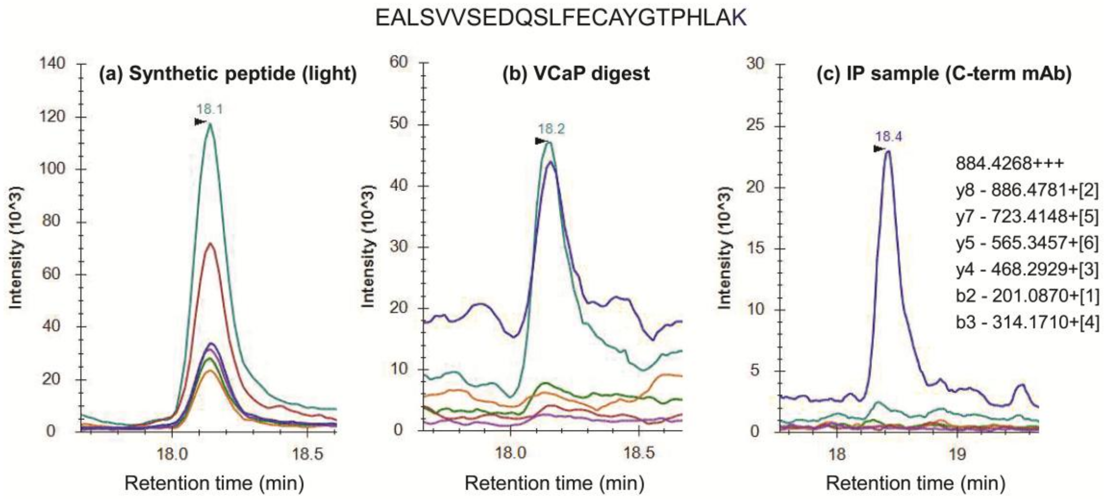


**Figure S2**. SRM measurement of the unique wild-type ERG peptide EALSVVSEDQSLFECAYGTPHLAK representing all ERG protein-coding isoforms, except A0A0C4DG41 (abundance 13.6 TMP; **Supplemental Table S1**), isoform-1 (12.1 TMP) and A0A088AWP2 (0.06 TPM). Chromatograms represent: (a) synthetic light peptide; (b) direct digest of VCaP lysate; and (c) ERG immunoprecipitated (IP) from the VCaP lysate with the C-term mAb. We thus conclude that two wild-type chromosomes 21 (lacking TMPRSS2-ERG gene fusions) of VCaP cells do not express any detectable amounts of the wild-type ERG.

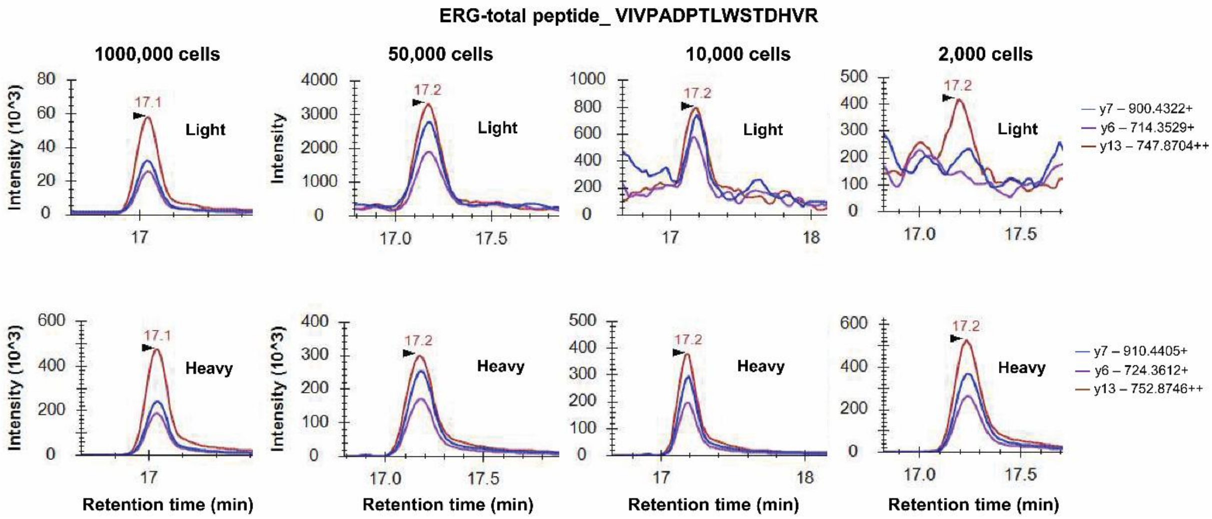


**Figure S3.** Limit of detection (LOD) of the IP-SRM assay using the total ERG peptide VIVPADPTLWSTDHVR was estimated at ~ 10,000 VCaP cell lysate equivalent.

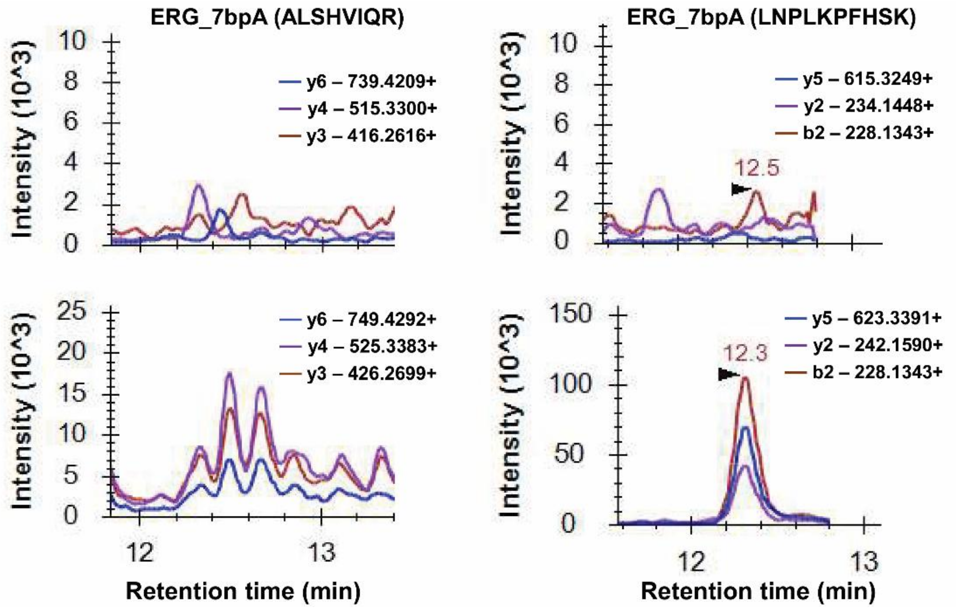


**Figure S4**. IP-SRM measurements of the endogenous peptides ALSHVIQR and LNPLKPFHSK representing ERG_7bpA protein isoform (isoform-8 mRNA). Peptide ALSHVIQR displayed a poor performance with significant interferences. No endogenous peptide LNPLKPFHSK of ERG_7bpA protein isoform was detected in VCaP cells.

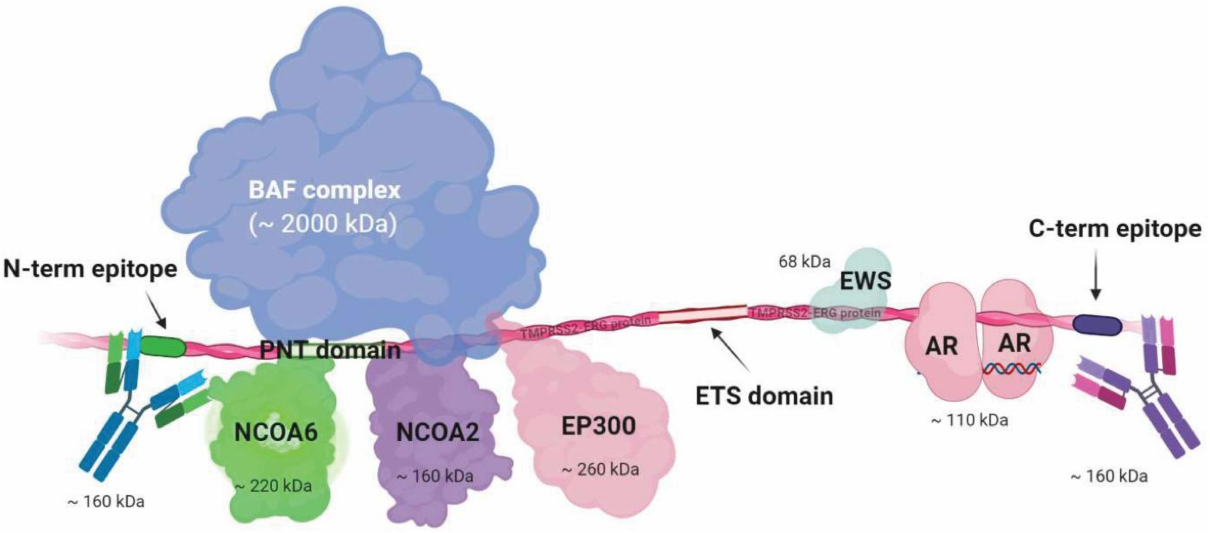


**Figure S5. Hypothetical illustration of the binding pattern of the identified ERG interactome, based on our and literature data.** The N-term mAb may interfere with the binding of very large proteins or complexes (BAF complex, NCOA2, NCOA6, and EP300 protein), while the C-term mAb does not interfere with the smaller EWS protein. Our results regarding AR binding were not consistent between the discovery (observed interaction with ERG) and independent verification experiments.

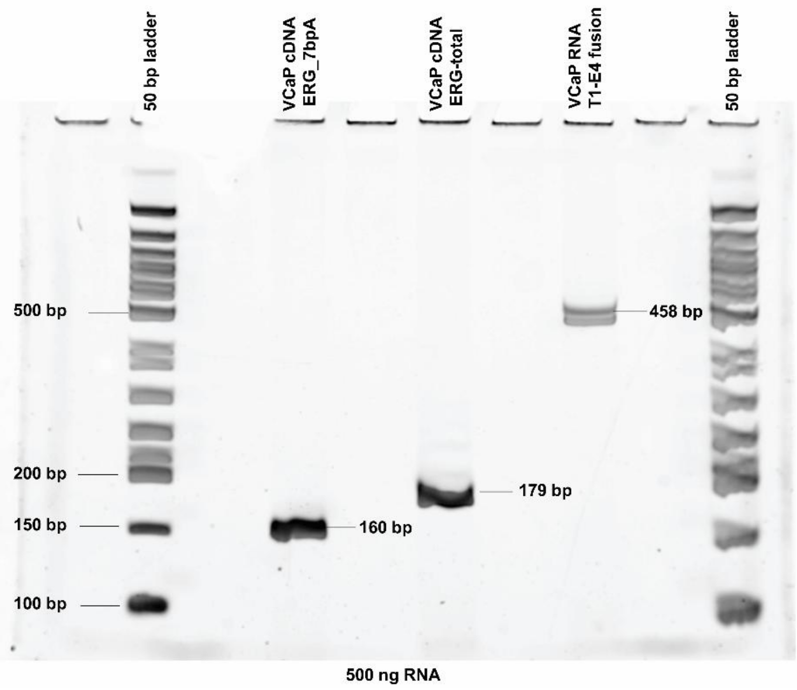


**Figure S6**. Validation of the presence of ERG_7bpA (160 bp) and ERG-total mRNA (179 bp) in VCaP cells by RT-PCR and agarose gel electrophoresis. The T1-E4 fusion (458 bp) was also included as a positive control. 500 ng RNA was used for RT-PCR.

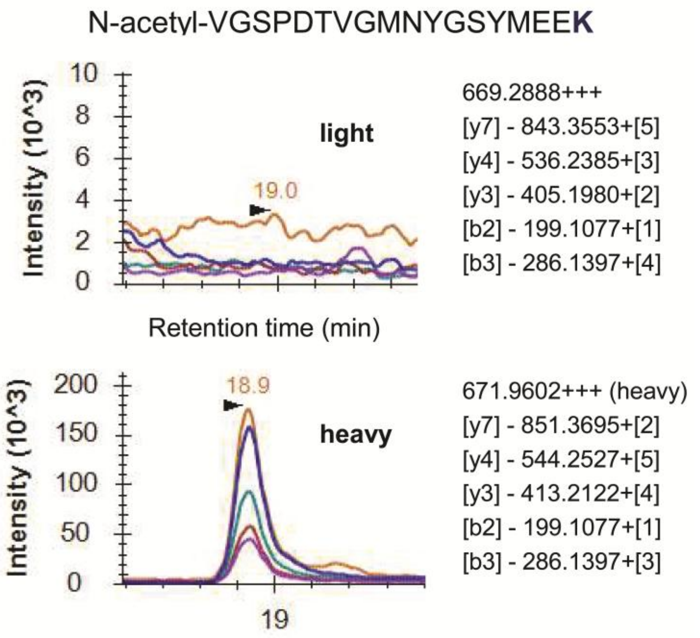


**Figure S7**. SRM measurement of the methionine-truncated and N-acetylated peptide VGSPDTVGMNYGSYMEEK for T1E4-ERG_Δ4 (14 % of the total ERG) and T1E4-ERG_Δ4Δ7b (1% of the total ERG) isoforms. ERG isoforms were immunoprecipitated from the VCaP lysate with the C-term mAb.
